# Transposon insertional mutagenesis in *Saccharomyces uvarum* reveals *trans*-acting effects influencing species-dependent essential genes

**DOI:** 10.1101/218305

**Authors:** Monica R. Sanchez, Celia Payen, Frances Cheong, Blake T. Hovde, Sarah Bissonnette, Adam P. Arkin, Jeffrey M. Skerker, Rachel B. Brem, Amy A. Caudy, Maitreya J. Dunham

## Abstract

To understand how complex genetic networks perform and regulate diverse cellular processes, the function of each individual component must be defined. Comprehensive phenotypic studies of mutant alleles have been successful in model organisms in determining what processes depend on the normal function of a gene. These results are often ported to newly sequenced genomes by using sequence homology. However, sequence similarity does not always mean identical function or phenotype, suggesting that new methods are required to functionally annotate newly sequenced species. We have implemented comparative analysis by high-throughput experimental testing of gene dispensability in *Saccharomyces uvarum*, a sister species of *S. cerevisiae.* We created haploid and heterozygous diploid Tn7 insertional mutagenesis libraries in *S. uvarum* to identify species dependent essential genes, with the goal of detecting genes with divergent functions and/or different genetic interactions. Comprehensive gene dispensability comparisons with *S. cerevisiae* predicted diverged dispensability at 12% of conserved orthologs, and validation experiments confirmed 22 differentially essential genes. Surprisingly, despite their differences in essentiality, these genes were capable of cross-species complementation, demonstrating that *trans-*acting factors that are background-dependent contribute to differential gene essentiality. This study demonstrates that direct experimental testing of gene disruption phenotypes across species can inform comparative genomic analyses and improve gene annotation. Our method can be widely applied in microorganisms to further our understanding of genome evolution.

## Introduction

The ability to accurately predict gene function based on DNA sequence similarity is a valuable tool, especially in the current stage of genomic research where an increasing number of genomes are being sequenced. It has become crucially important to predict gene function based on sequence similarity due to the lack of experimentally determined functional information associated with each newly sequenced genome. Most functional predictive methods rely on similarities of DNA sequence homology, co-expression patterns, or protein structure to help assign function to uncharacterized genes, using genes where known functions have been previously characterized (Eisen 1998; Usadel et al. 2009). However, these methods come with their own set of limitations and often produce a substantial number of predictive errors, highlighting the importance of implementing experimental methods to directly test gene function in previously uncharacterized genomes to improve current methods of annotation.

The gold standard of gene function characterization relies on observing phenotypes of targeted deletions of predicted coding sequences to probe the contributions of each gene to specific biological processes. To get a global view of gene function within an organism, several genome-wide deletion collections have been created in model species, particularly in bacteria and yeast (Baba et al. 2006; Berardinis et al. 2008; Porwollik et al. 2014; Winzeler et al. 1999), including highly diverged species (Kim et al. 2010; Schwarzmüller et al. 2014) as well as different strains within a species (Dowell et al. 2010). These systematic deletion collections are powerful tools for investigating gene function, biological pathways, and genetic interactions, especially in the genetic workhorse *Saccharomyces cerevisiae,* where gene function characterization and gene dispensability comparisons have been extensively performed amongst various deletion collections of yeast (Costanzo 2016; Dowell et al. 2010; Kim et al. 2010; Tong et al. 2001). These studies have identified approximately 17% of essential genes to be differentially essential between highly diverged species (*S. cerevisiae* and *Schizosaccharomyces pombe*) and have discovered 6% of essential genes (57) that are differentially essential even between two strains of *S. cerevisiae.*

However, considerable effort and resources are required to create these targeted, systematic libraries, and they are not a practical approach for interrogating a wide range of non-standard genetic backgrounds in a high-throughput manner. Alternative approaches to targeted gene deletion libraries are transposon-based mutagenesis methods used to create random insertional mutant collections, eliminating requirements for *a priori* knowledge about defined coding regions and providing information about partial loss-of-function or gain-of-function mutations. Random insertional profiling has been widely applied across various species and has been instrumental in understanding virulence genes, stress tolerance mechanisms, and even tumor suppressor genes in mice (DeNicola et al. 2015; van Opijnen and Camilli 2013; de la Rosa et al. 2017; Weerdenburg et al. 2015; Yung et al. 2015; Coradetti et al. 2018). In yeasts in particular, transposon libraries have provided useful information about gene function, growth inhibiting compounds, and essential functional protein domains (Gangadharan et al. 2010; Guo et al. 2013b; Michel et al. 2017; Oh et al. 2010; Ross-Macdonald et al. 1999; Zhao et al. 2017; Price et al. 2018; Zhu et al. 2018).

Despite this growing body of literature, the genetic mechanisms explaining differences across species are still poorly understood. Here we utilize a random insertional method to assay gene dispensability using ~50,000 mutants in *Saccharomyces uvarum,* a species that diverged from *S. cerevisiae* 20 million years ago and whose coding sequences are ~20% divergent from those of *S. cerevisiae* (Dujon 2010; Kellis et al. 2003; Scannell et al. 2007). These species can mate with one another to create hybrids, allowing us to explore the genetic basis for possible differential gene dispensability between them using genetic toolsets established in *S. cerevisiae.* Genes with different dispensability patterns between these two species could be explained by divergent gene function and/or genetic interactions, providing a model for investigating genome evolution between two diverged species of yeast. We successfully validate a subset of predicted differentially essential genes required for growth in rich media, establishing the utility of our mutagenesis approach in prioritizing genes for testing viability (Michel et al. 2017; Guo et al. 2013a). In what follows, for genes that emerge from our analyses as differentially essential between the reference strains of *S. cerevisiae* and *S. uvarum*, we refer to them as species-specific; rigorously speaking, a comprehensive study across populations will be necessary to establish whether a given essentiality pattern is truly common to the entirety of the respective species. Together, our data make clear that our Tn7 transposon mutagenesis library serves as a valuable resource for studying the *S. uvarum* genome, and that our approach is a powerful framework for comparative functional genomics studies across newly sequenced, previously uncharacterized species.

## Results

### Generating Tn7 insertional libraries in *S. uvarum* to predict essential and nonessential genes

One of the most straightforward mutant phenotypes to characterize is cell viability, which reveals if a given gene is involved in an essential cellular process. Therefore, we first sought to characterize gene essentiality in *S. uvarum*, with the aim of identifying genes that are differentially essential between *S. cerevisiae* and *S. uvarum*. Instead of creating a library of individual knock-out strains, we applied a high-throughput approach of creating random insertional mutants and leveraged the power of sequencing to identify the insertion sites in a pooled collection. The Tn7 mutagenesis library approach described by Kumar *et al.,* (2004) was used to create a collection of *S. uvarum* mutant strains and has been previously described by Caudy *et al.,* (2013). Briefly, *in vitro* transposition of the Tn7 transposon was performed in a plasmid library containing random *S. uvarum* genomic fragments. The Tn7 transposon was designed to carry a ClonNat resistance marker that carries stop codons in all reading frames near both termini. The interrupted genomic fragments were excised out of the plasmid and integrated at their corresponding genomic positions in the reference strain background of *S. uvarum*, each of which is expected to produce a truncation when inserted within a coding region (**Figure 1A**). The plasmid library contains ~50,000 unique genomic insertion sites; we integrated the library into a diploid strain and, separately, a haploid *MAT*a strain at 10X coverage (additional details can be found in **Supplemental Information**). Pools of mutants from each Tn7 library were grown in liquid nutrient rich media as described in Materials and Methods. Insertion sites were determined using sequencing methods as described in detail in the **Supplemental Information** along with DNA sequencing library preparation protocols (**Figure 1C**).

**Figure 1.**
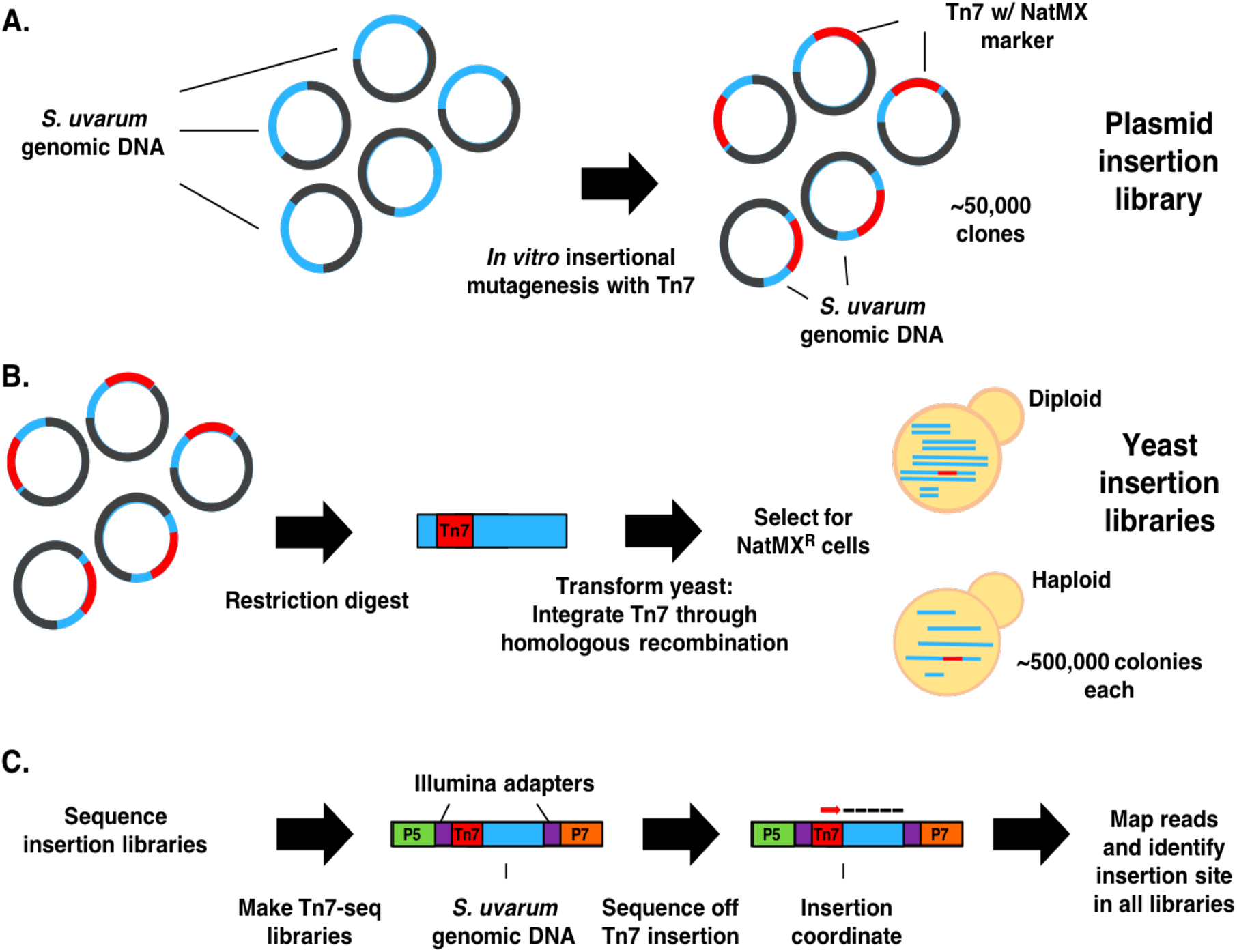
Schematic of Tn7 transposon mutagenesis library and insertion identification in *S. uvarum.* **A)** Simplified representation of *in vitro* transposition of the Tn7 transposon into a plasmid library containing random *S. uvarum* genomic DNA fragments. Approximately 50,000 plasmids containing the Tn7 transposon were pooled together to form the final library. **B)** Illustration of the Tn7 containing excised portion of the plasmid integrated into haploid and diploid yeast through homologous recombination. Approximately 500,000 Nat^R^ clones of each ploidy were pooled into two separate pools (Haploid pool and Diploid pool). **C)** Design of Tn7-seq libraries used to identify insertion sites through sequencing. Reads containing Tn7 sequence are enriched (PCR off common flanking region of the Tn7 and Illumina adapter sequences) and mapped to the genome to identify insertion sites.

We catalogued transposon insertion mutants on the basis of sequenced insertion sites that could be detected after mutagenesis and outgrowth of haploid and diploid *S. uvarum*, and we also tabulated those mutants present in both pools. We found these insertion sites to be evenly distributed throughout the *S. uvarum* genome, as illustrated in **Supplemental Fig. 1**. (Detailed information about overall sequencing coverage is listed in **Supplemental File 4.**) Once the insertion sites were determined in both libraries, we counted the number of insertion sites in each annotated open reading frame (Materials and Methods). **Supplemental Table 1** summarizes the number of insertion sites and the number of genes that contain insertion sites within each library, including the initial plasmid library. The number of insertion sites from each library that fell into each annotated *S. uvarum* gene is listed in **Supplemental File 5.** Of the 5,908 annotated genes, a total of 5,315 (90%) genes harbored insertion sites that were identified in at least one library. Comparisons between shared genes and unique genes harboring insertion sites are illustrated in **Supplemental Fig. 2**.

Since by definition, loss of function mutants in genes essential in *S. uvarum* would not be viable in the haploid of this species, we used our observations of detected transposon mutants in this strain as a jumping-off point for inferences of gene essentiality. We first tested whether orthologs of the known essential set in *S. cerevisiae* would be depleted for insertion sites in the *S. uvarum* haploid, as would be predicted if the essentiality of most genes were conserved between the species. Consistent with this notion, we identified a significant reduction in the number of inserts present in known *S. cerevisiae* essential genes in the haploid *S. uvarum* library (Wilcoxon test p < 2.2 e^−16^, essential average inserts/kb=0.88, SD=1.28 vs. non-essential average inserts/kb = 4, SD = 4.38) (**Supplemental Fig. 3**), indicating that essential genes are effectively targeted by this approach. However, due to the nature of the library, insertional events at different positions across a gene may result in a partial loss-of-function (Sadhu et al. 2018). Since essential genes may still tolerate some insertions, we instead relied on comparisons between the diploid and haploid libraries to make inferences about gene essentiality. Specifically, we calculated an insertion ratio using the number of inserts per gene in the haploid library divided by the number of inserts in the diploid library, which inherently normalizes for the length of the gene (Materials and Methods). Using the insertion ratio as a metric, we tested for significant differences between *S. uvarum* genes whose orthologs were essential and non-essential in *S. cerevisiae.* In our analyses, we used *S. uvarum* intergenic regions as a control: intergenic regions between convergently oriented genes are expected to largely not be essential, and thus we expected the distribution of intergenic regions to be similar to that of non-essential genes.

**Figure 2A** reports the distribution of detected insertion mutants in *S. uvarum* haploids and diploids (quantified by the insertion ratio) for each feature type. As predicted, we found that *S. uvarum* orthologs of known *S. cerevisiae* non-essential genes generally behaved similarly in our mutant pools to *S. uvarum* intergenic regions. Interestingly, however, the distribution of insertion ratios from these orthologs of non-essential genes had a left shoulder resembling the distribution among orthologs of *S. cerevisiae* essential genes. We hypothesized that this population of genes depleted for insertions in *S. uvarum* haploids was likely to reflect *S. uvarum-*specific essential genes. Likewise, we also noted a right-hand tail (corresponding to highly abundant transposon mutants in *S. uvarum* haploids) in the distribution of insertion ratios among *S. uvarum* orthologs of *S. cerevisiae* essential genes, suggesting that some were in fact not essential in *S. uvarum*. The differences between orthologs of *S. cerevisiae* essential genes and non-essential gene insertion ratios were significant, as well the differences between orthologs of essential genes and intergenic regions (**Figure 2B**, Wilcoxon p < 2.2e^−16^). Using the insertion ratio, we formulated a prediction of each gene as either essential or non-essential in *S. uvarum* using a null distribution to rank genes above or below a cut-off metric of 0.25 (details described in **Supplemental Information**). Using this cut-off value, 1170 genes were categorized as predicted essential genes. We applied an additional cut-off metric (more details in Material and Methods) to remove a class of low coverage genes, resulting in a total number of 718 (13%) predicted essential genes and 3,838 (65%) genes that are predicted non-essential, with 1299 genes (22%) undetermined (genes without inserts in the diploid library). We proceeded to characterize each gene set and validate the dispensability of each of the predicted gene categories.

**Figure 2.**
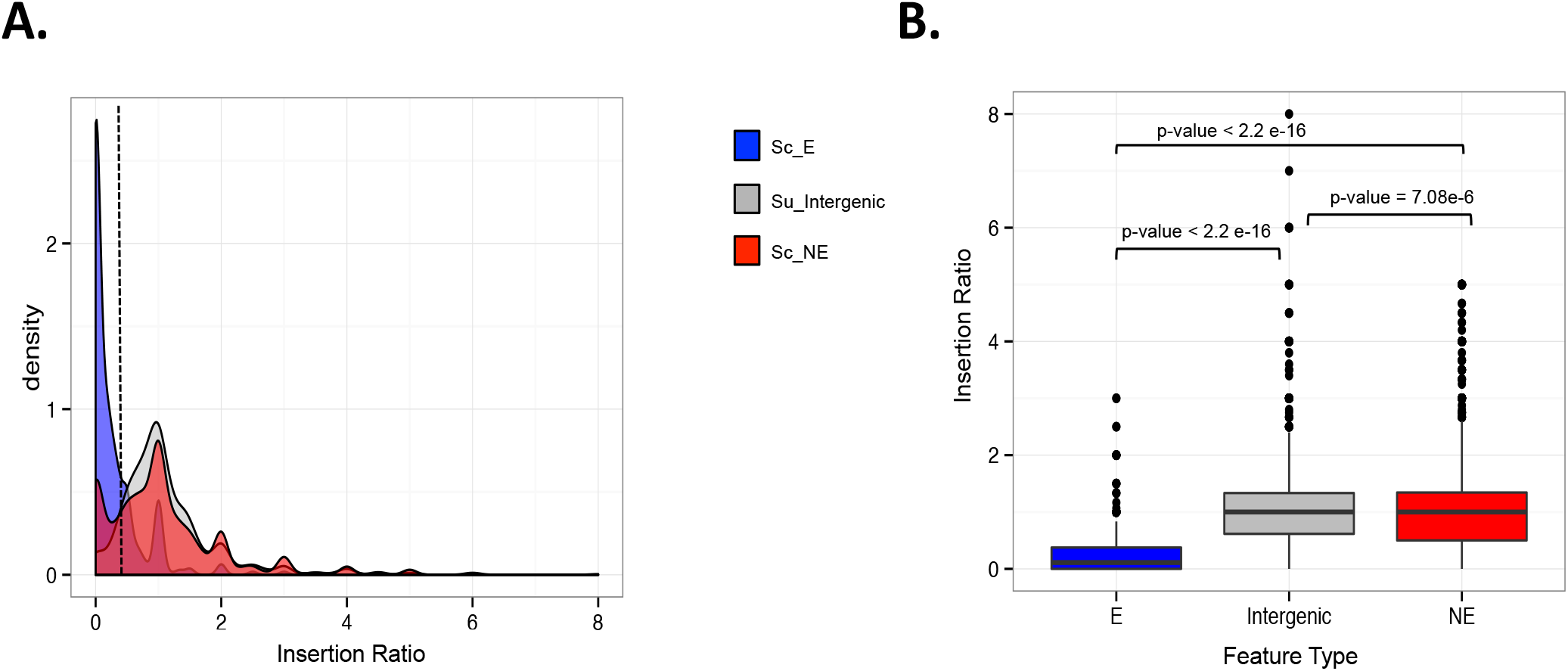
Insertion ratio distributions of *S. uvarum* intergenic regions and known *S. cerevisiae* essential and non-essential genes. **A)** Density plots displaying the distribution of insertion ratios across three feature types: *S. uvarum* intergenic regions between Watson and Crick oriented genes ranging from 7 kb-500 bp (grey) and *S. uvarum* genes whose orthologs are known *S. cerevisiae* essential (Sc_E in blue) and nonessential genes (Sc_NE in red). The dashed line represents an insertion ratio of 0.25 and defines the cut-off value to classify essential and non-essential genes. **B)** Box plots of insertion ratios by feature type described in plot A. Significant insertion ratio differences exist between known *S. cerevisiae* essential and non-essential genes and between *S. uvarum* intergenic regions. (Wilcoxon tests Sc_E:Su_Intergenic p < 2.2e^−16^, Sc_E:Sc_NE p < 2.2e^−16^, Sc_NE:Su_Intergenic p = 7.08e^−6^).

### Analysis of predicted gene dispensability

The predicted gene list of *S. uvarum* essential genes was compared to known essential genes lists from both *S. cerevisiae* and *S. pombe* to determine the amount of conservation that exists between orthologs across diverged species. Of the predicted 718 *S. uvarum* essential genes, 297 genes (42%) were shared amongst all three sets, with a total of 487 genes (68%) shared with at least one other set. Furthermore, 9 genes whose essentiality was specific to particular *S. cerevisiae* strains, including 4 genes specific to the S288C strain and 5 genes that are specific to the ∑1278b strain, were inferred by our analysis to be essential in *S. uvarum* (**Supplemental Fig. 4**). Similar to what has been previously shown in *S. cerevisiae,* predicted essential genes in *S. uvarum* were more likely to be unique, with 91% of essential genes (656/718) being present in single copy compared to 76% of non-essential genes (2736/3604). Additionally, comparisons between Gene Ontology (GO) molecular function terms of essential gene sets from both species showed significant enrichment (p-value < 0.01) for fundamental biological functions. Processes such as DNA replication/binding, RNA and protein biosynthesis, as well as structural constituents of the ribosome and cytoskeleton were enriched in both predicted *S. uvarum* essential genes and those known to be essential in *S. cerevisiae* (**Supplemental File 6**). In contrast, non-essential genes were significantly (p-value < 0.01) enriched for regulatory functions (transcription factor activity) and conditional responsive processes, such as transmembrane transporter activity and cell signaling (kinase activity) (**Supplemental File 7**). We conclude that many of the features of the predicted essential genes in *S. uvarum* are similar to confirmed essential genes in other species.

We next sought to validate experimentally the predictions of essentiality we had made from our transposon mutagenesis. We first focused on genes whose orthologs were known to be essential in *S. cerevisiae*, and which we had likewise predicted to be essential in *S. uvarum*. For each of 13 such cases, we sporulated the respective heterozygous deletion strain and performed tetrad analysis for cell viability; the results confirmed essentiality for 12 (92%) of the 13 strains (**Supplemental Table 2**). One example of a confirmed essential gene can be found in **Figure 3A**, which illustrates the genomic positions of all insertion sites across a genomic locus of chromosome V that contains essential and non-essential genes. The color of the gene outline reports the predicted dispensability, which is determined by their insertion ratio. For example, the gene *BRR2,* a RNA-dependent RTPase RNA helicase, had an insertion ratio of 0.130 and was predicted to be a conserved essential gene (**Figure 3C**). The tetrad analysis of a *BRR2* heterozygous deletion strain displayed a 2 viable:2 inviable segregation pattern in both *S. cerevisiae* and *S. uvarum*, validating this gene as a conserved essential gene (**Figure 3B**). Images of all other confirmed essential genes are in **Supplemental Fig. 5.** We also tested three genes known to be non-essential in *S. cerevisiae* and predicted by our analysis to be non-essential in *S. uvarum*, confirming all three (100%) as non-essential in both species (**Supplemental Fig. 6**) (**Supplemental Table 2**).

**Figure 3.**
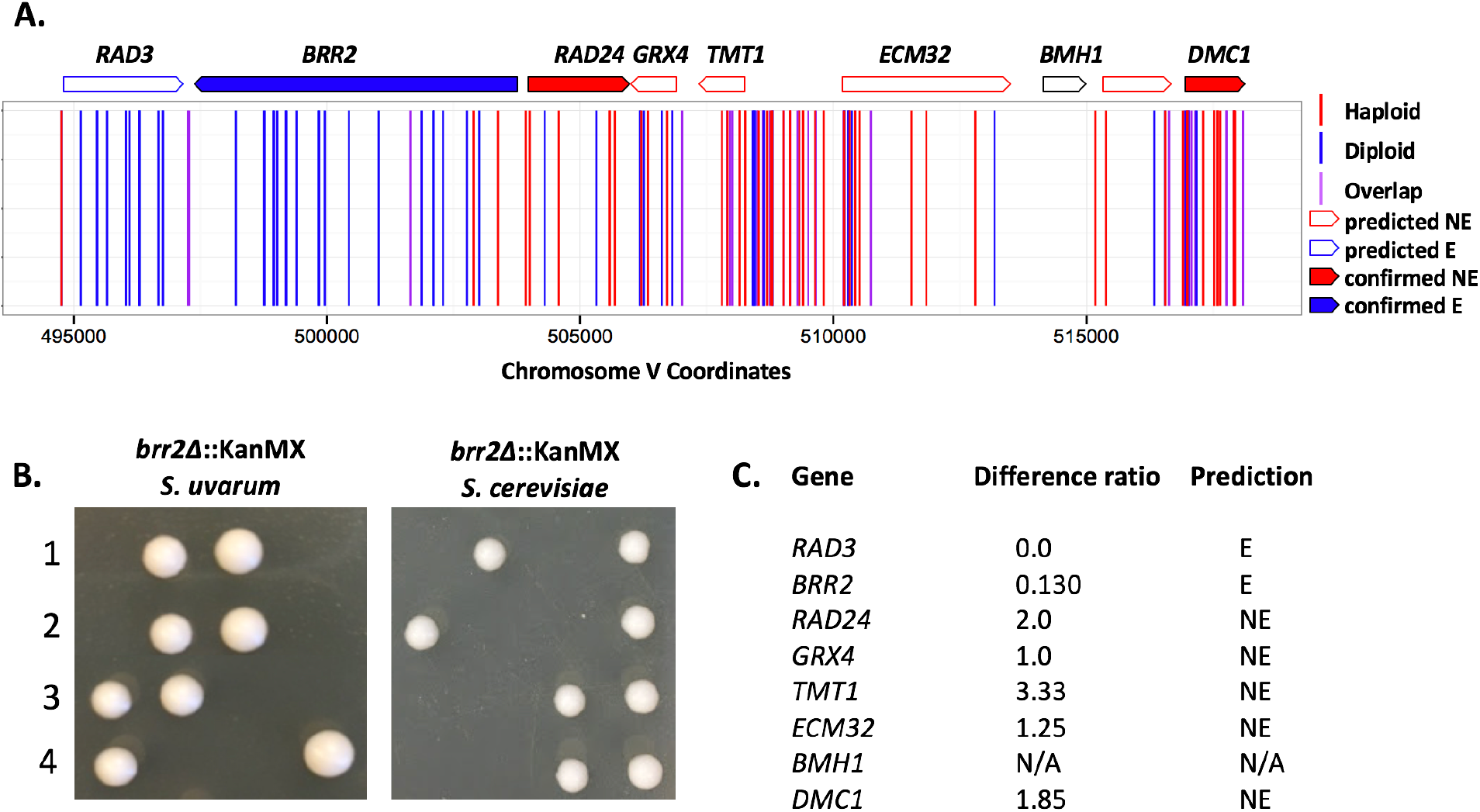
Validation of conserved essential and non-essential genes. A) Mapped chromosomal insertion positions are plotted across chromosome V. Haploid inserts are indicated in red, diploid inserts are blue, and overlapping inserts are indicated in purple. Genes indicated across the top are outlined according to predicted dispensability and filled in if confirmed. B) Tetrad analysis of a confirmed conserved essential gene *brr2∆* in *S. cerevisiae* and *S. uvarum.* Segregants containing *brr2∆* alleles are inviable in both species. C) Table indicating the insertion ratio (number of haploid inserts by the number of diploid inserts) per gene. The final column lists the predicted classification (NE, non-essential; E, essential; N/A, no data).

Additionally, we obtained an independent set of *S. uvarum* haploid deletion strains (see Materials and Methods), which was used as a validated non-essential gene set. Out of the total 356 genes that were viable upon deletion in this collection, our analysis of transposon mutants inferred 346 to be non-essential in *S. uvarum* (97%), providing further strong support to the predictions from our method.

### Gene dispensability comparisons of orthologous pairs between *S. cerevisiae* and *S. uvarum*

Our main goal of this project was to identify genes that were differentially essential in a species-dependent manner. To make direct comparisons of dispensability between *S. cerevisiae* and *S. uvarum,* we narrowed our analysis to 4,543 orthologous genes for which we had data in the *S. uvarum* dataset (**Supplemental File 8**). Overall, our predicted patterns of essentiality in *S. uvarum* were consonant with the known behavior in *S. cerevisiae* for 88% (4016/4543) of these genes. The remaining 12% of orthologs were predicted to differ in essentiality between the two species, with 304 (7%) of these genes only essential in *S. uvarum* and 221 (5%) genes only essential in *S. cerevisiae* (**Supplemental Figure 7A**). We note that the former could represent an over-estimate of the count of predicted essential genes specific to *S. uvarum,* in that it is inferred from the lack of detected insertion sites in our haploid *S. uvarum* libraries in genes not previously characterized as essential in *S. cerevisiae*. All predicted genes that differ in dispensability are listed in **Supplemental File 9.**

To analyze further our predictions of differential essentiality between the species, we compiled a list of 222 genes whose respective orthologs were known to be essential in *S. cerevisiae* and predicted to be non-essential in *S. uvarum*; we also formulated more stringent cutoffs for our transposon mutant library analysis (see Materials and Methods) to yield a similarly sized set of genes (220 total) inferred to be essential in *S. uvarum* and known to be non-essential in *S. cerevisiae*. Using this list, we determined the proportion of inferred *S. cerevisiae-*specific and *S. uvarum-*specific essential genes annotated for each function by performing Gene Ontology (GO) term finder using the molecular function ontology. Among the results, reported in **Supplemental Figure 7B,** most striking were enrichments for structural constituents of the ribosome among genes predicted to be essential only in *S. uvarum,* and for RNA polymerase activity among genes inferred to be essential only in *S. cerevisiae*. Full lists of significant (p-value < 0.01) GO enrichment molecular function terms for each species individually are listed in **Supplemental File 10.** We also hypothesized that genetic interaction patterns could distinguish genes that were predicted to be differentially essential between *S. cerevisiae* and *S. uvarum*. Toward this end, we tabulated combined interaction degree scores for all orthologous genes between yeast, worm, flies, mice, and humans. We first compared interaction degrees between known essential and non-essential genes, and found the former to be increased, as previously reported (Costanzo 2016) (**Supplemental Fig. 8**). Next, we formulated a comparison between two gene sets: those predicted to be essential in *S. uvarum* and known to be non-essential in *S. cerevisiae* on the one hand, and those categorized as non-essential in both species on the other hand. We found a striking and significant increase in the number of combined interactions in the former (p-value = 1.99 e^−7^), suggesting that having more interactions may be predictive of essential function. Analyses of expression could not account for the differences in essentiality (**Supplemental Figure 13**).

We next set out to confirm experimentally a subset of predicted essential genes within each species, by sporulating heterozygous deletion strains to determine the viability pattern of the segregants. As an example of a confirmed *S. uvarum-*specific essential gene from these experiments*, SSQ1,* which is required for assembly of iron/sulfur clusters into proteins, is illustrated in **Figure 4**. **Figure 5** displays an example of a gene confirmed to be essential in *S. cerevisiae* but not *S. uvarum*: *VTC4,* a gene involved in the regulation of membrane trafficking. Overall, we confirmed a total of 22 *S. uvarum-*specific and *S. cerevisiae*-specific essential genes (tetrad analysis can be found in **Supplemental Figs. 9 and 10** respectively). Interestingly, we found a variety of growth phenotypes associated with deletion in the permissive species background of confirmed species-specific essential genes, with some deletions showing poor growth and others growing as well as wt. All combined tetrad analysis results are reported in **Supplemental Fig. 11**; these results include genes for which sporulation experiments did not validate the inferences of species-specific essentiality from our transposon mutagenesis, which we refer to as false positives. **Supplemental Table 2** summarizes the total number of genes confirmed in each category. We note the higher false positive rate in the species-specific essential gene categories; as above, in part this likely reflects incorrect calls of essentiality in *S. uvarum* in which we detected no transposon mutants in our haploid libraries as a product of low coverage, rather than inviability of the respective mutants.

**Figure 4.**
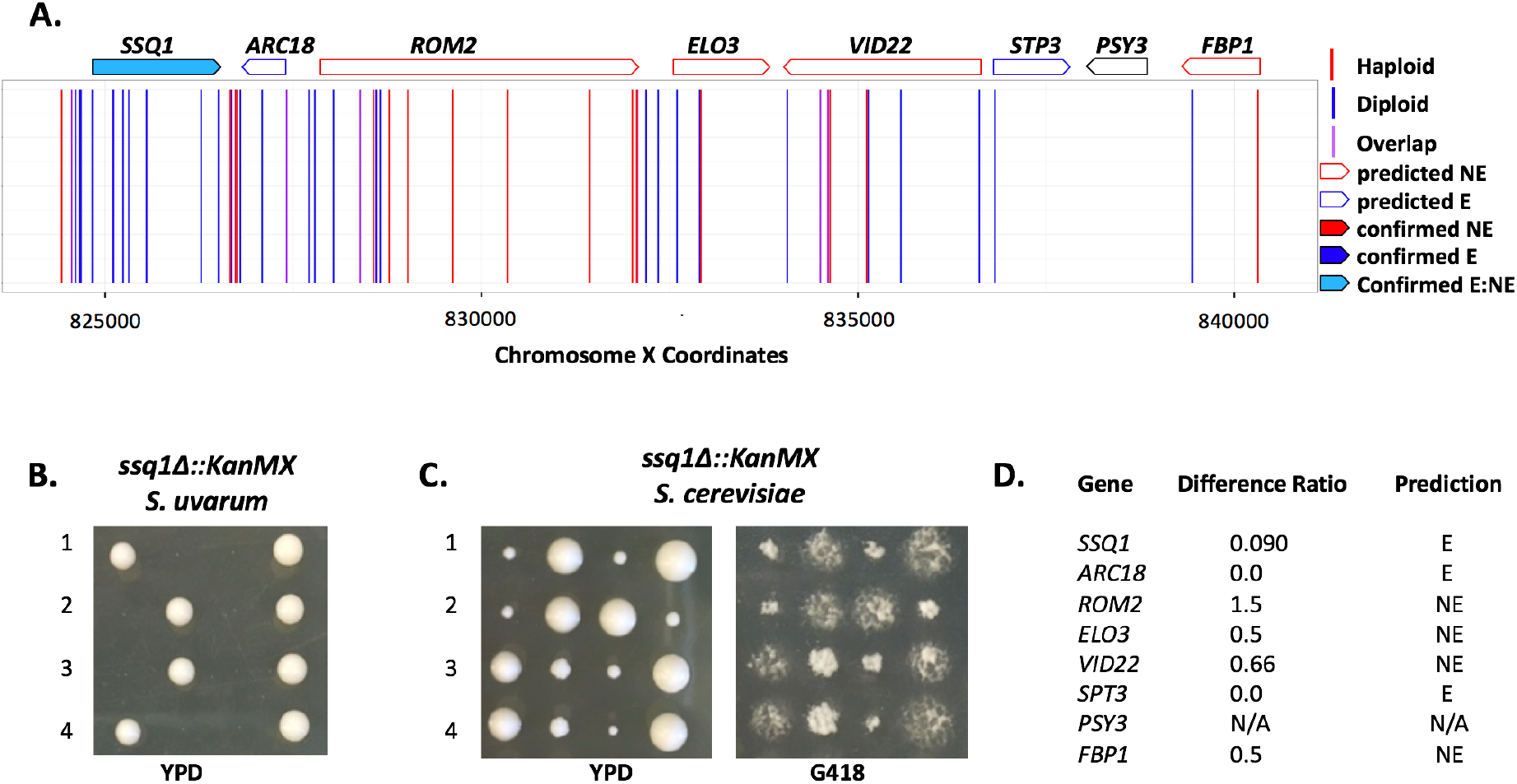
Validation of *S. uvarum-*specific essential gene *SSQ1*. A) Mapped chromosomal insertion positions are plotted across chromosome X. Haploid inserts are indicated in red, diploid inserts are blue and overlapping inserts are indicated in purple. Genes indicated across the top are outlined according to predicted dispensability and filled in, if confirmed. Light blue filling indicates a gene that is essential in *S. uvarum* and non-essential in *S. cerevisiae* (confirmed E_NE). B) Tetrad analysis of a heterozygous *ssq1∆::KanMX* strain displaying inviable segregants containing the *ssq1∆* allele in *S. uvarum.* C) Tetrad analysis of a heterozygous *ssq1∆::KanMX* strain in *S. cerevisiae* containing viable segregants plated on YPD and G418. D) Table indicating the insertion ratio (number of haploid inserts by the number of diploid inserts) per gene. The final column lists the predicted classification (NE, non-essential; E, essential; N/A, no data).

**Figure 5.**
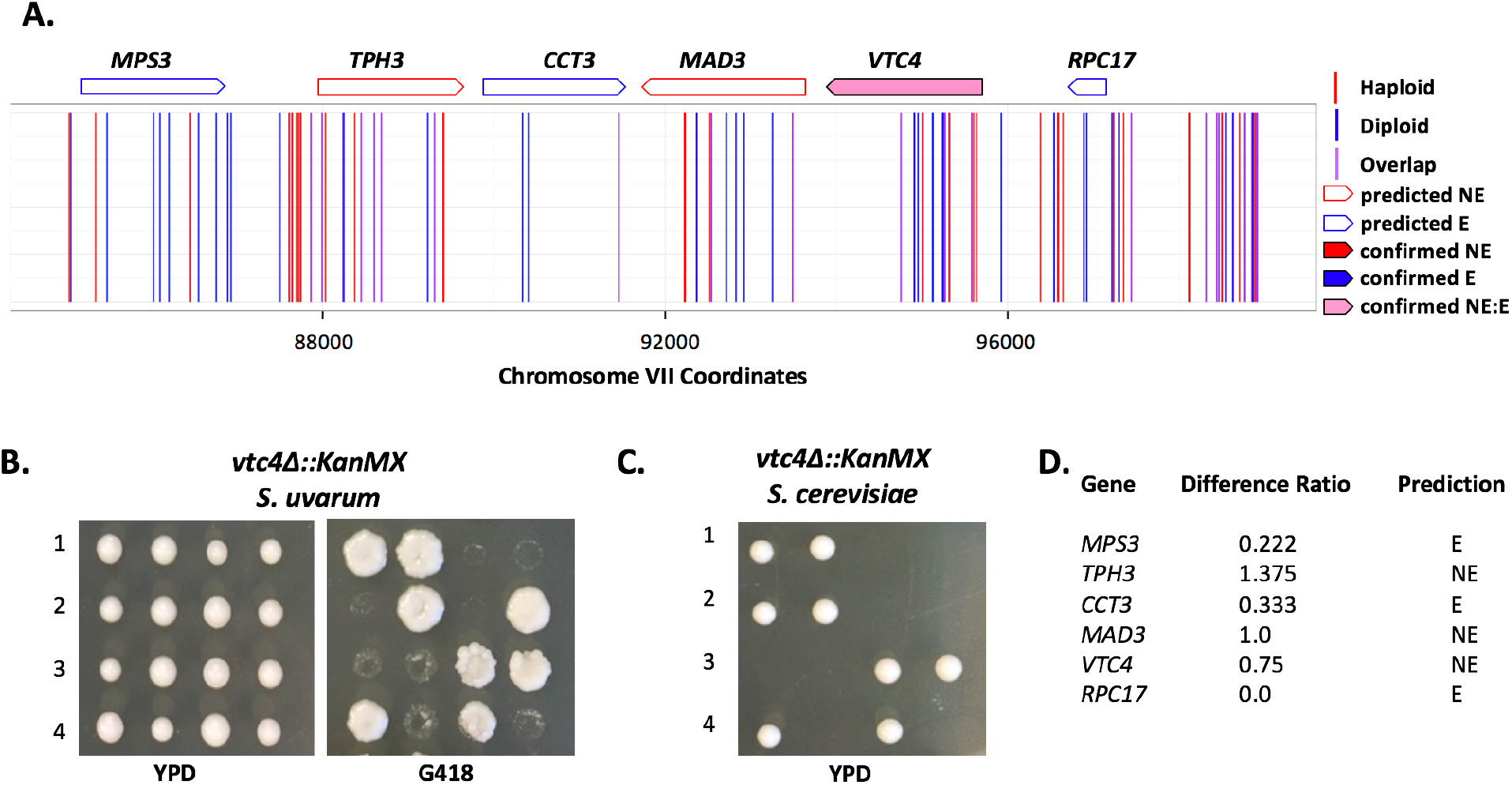
Validation of *S. cerevisiae-*specific essential gene *VTC4*. A) Mapped chromosomal insertion positions are plotted across chromosome XII. Haploid inserts are indicated in red, diploid inserts are blue and overlapping inserts are indicated in purple. Genes indicated across the top are outlined according to predicted dispensability and filled in if confirmed. Light pink filling indicates a gene that is essential in *S. cerevisiae* and non-essential in *S. uvarum* (confirmed NE_E). B) Tetrad analysis of a heterozygous *vtc4∆* strain displaying viable segregants with the *vtc4∆* allele in *S. uvarum* plated on YPD and G418. C) Tetrad analysis of a heterozygous *vtc4∆* allele in *S. cerevisiae* resulting in inviable segregants. D) Table indicating the insertion ratio (number of haploid inserts by the number of diploid inserts) per gene. The final column lists the predicted classification (NE, non-essential; E, essential; N/A, no data).

That said, in some cases, validation failures could be explained by errors in genome annotation, not errors in our insertional library results. For example, the gene *DRE2,* which functions in cytosolic iron-sulfur protein biogenesis, was predicted to be a non-essential gene, but was confirmed as an essential gene through tetrad analysis. Manual inspection revealed that all the haploid insertions were clustered at the 5’ end of the gene (**Supplemental Fig. 12**). In protein alignments of *DRE2* between *S. cerevisiae* and *S. uvarum,* we noted an annotated start codon in *S. uvarum* upstream of the annotated start codon in *S. cerevisiae.* These data strongly suggest that the gene was misannotated in *S. uvarum*, and instead shares the methionine start position further downstream. Using the reannotated gene coordinates, we would correctly classify *DRE2* as essential in *S. uvarum* since the haploid insertions were no longer included in the open reading frame. This example highlights the utility of our method to improve gene annotation in addition to characterizing gene essentiality.

### Paralog divergence and duplicate gene loss explain some background effects on differential gene essentiality

We next set out to investigate genetic background effects that could be contributing to differences in gene dispensability between *S. cerevisiae* and *S. uvarum.* One explanation could be genetic redundancy due to gene duplications, such that a gene is nonessential in one species due to the presence of a paralog, whereas the other species contains only a single copy. To investigate this possibility, we began investigating genes that both differed in dispensability and harbored a paralog. Of the 222 *S. cerevisiae-*specific essential genes, 11 were known to have paralogs. Our initial analysis identified the Ras activator *CDC25* as a *S. cerevisiae-*specific essential gene (nonessential in *S. uvarum*)*. CDC25* is a paralog of *SDC25*, which contains a premature stop codon in the reference strain of *S. cerevisiae* and other laboratory strains (Folch-Mallol et al. 2004).

We performed complementation assays by cloning *S. uvarum* alleles of both paralogs into a CEN/ARS plasmid and testing whether the *S. uvarum* alleles could rescue the inviable phenotype of segregants from a heterozygous *cdc25∆* deletion in *S. cerevisiae*. We found that *SDC25* from *S. uvarum* was functional, and that both *SDC25* and *CDC25* alleles from *S. uvarum* could complement a *cdc25∆* deletion in *S. cerevisiae* (**Table 1**). Although we did not test for complementation in the *S. uvarum* background, the results from the complementation assays demonstrate that *CDC25* is required for growth in this strain of *S. cerevisiae* due to the lack of redundancy as a consequence of the non-functional copy of *SDC25*. We expect that this relationship may be unique to those *S. cerevisiae* strains in which *SDC25* is a pseudogene, but we nonetheless consider it a satisfying validated mechanism for the background dependence of *CDC25* essentiality.

**Table 1.** Viability summary of gene deletions and complementation assays. Signs in columns indicated by each species represent viability (-:inviable, +:viable, N/D: not done). Complementation assays are represented by the gene deletion with the addition of each gene expressed on a low copy plasmid.

| Genotype | *S. cerevisiae* | *S. uvarum* |
| --- | --- | --- |
| *cdc25Δ* | - | + |
| *sdc25Δ* | + | - |
| *cdc25Δ + ScCDC25* | + | N/D |
| *cdc25Δ + SuCDC25* | + | N/D |
| *cdc25Δ + SuSDC25* | + | N/D |
| *cdc25Δ + ScSDC25* | - | N/D |
| *sdc25Δ + ScCDC25* | N/D | - |

Following this same logic, we hypothesized that *CDC25* non-essentiality in *S. uvarum* could be attributed to the redundancy provided by the functional copy of *SDC25* in this species. To test this idea, we created an *S. uvarum* mutant heterozygous for both *cdc25∆* and *sdc25∆* and performed segregation analysis on the dissected tetrads (**Table 1**). Unexpectedly, the segregation pattern of a double mutant displayed a lethal phenotype for not only the double mutant but also the single *sdc25∆* mutant. We confirmed this result by constructing an *sdc25∆* heterozygous mutant in *S. uvarum* and found a 2:2 segregation pattern showing that *SDC25* is an essential gene in *S. uvarum.* We conclude that, of the two paralogs, one is essential in our *S. cerevisiae* strain and the other in *S. uvarum*, representing a novel case of a swap in essentiality.

### Divergent gene dispensability is largely due to *trans* effects

While genetic redundancy or gene loss is a possible explanation for a fraction of differentially essential genes, the remaining much larger portion of this class of genes remained unexplained. Another hypothesis to explain differences in essentiality is gene function divergence between these two species. We therefore proceeded to further investigate the remaining differentially essential genes for functional differences. For a subset of these genes, we performed complementation assays in both species to test for divergent function. We cloned five *S. cerevisiae* alleles and their promoters from the list of *S. uvarum-*specific essential genes (*SAC3, TUP1, CCM1, SSQ1,* and *AFT1*) and seven *S. uvarum* alleles from the list of *S. cerevisiae-*specific essential genes (*ALR1*, *SHR3*, *CDC25*, *INN1*, *LCD1*, *SEC24*, *VTC24*) and tested each allele’s ability to rescue the inviability caused by deletion of the corresponding ortholog in the alternate species (**Figure 6**). The results from these complementation tests revealed that all genes are able to complement the inviable phenotype in the other species, suggesting that the differences in essentiality are more likely to be due to *trans*-acting changes rather than functional differences of protein-coding regions.

**Figure 6.**
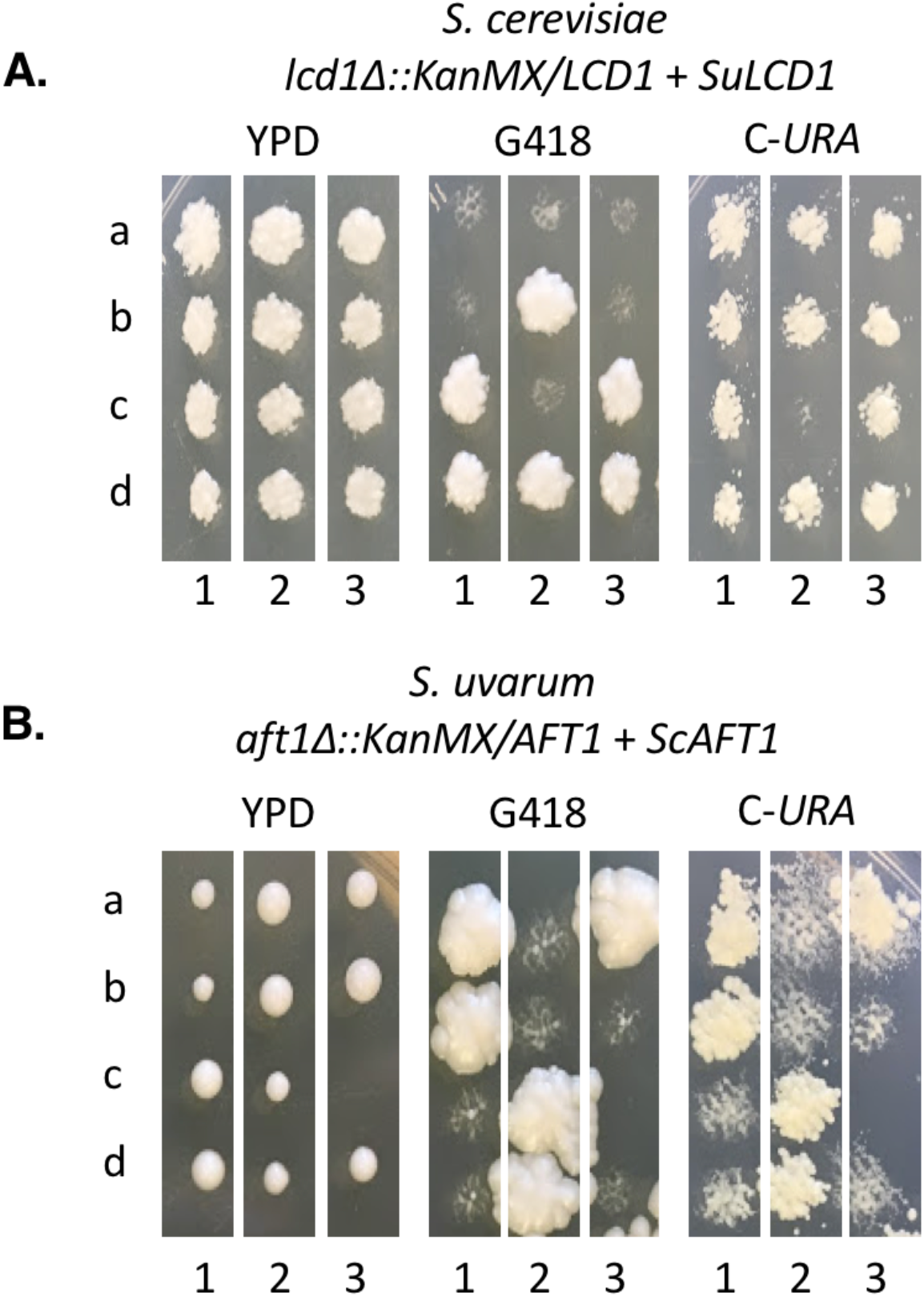
Complementation assay confirming two examples of genes that differ in essentiality but complement the viability phenotype in both genetic backgrounds. Letters a-d represent four spores, while numbers 1-3 indicate three tetrads. Tetrads were dissected on YPD and were replica plated to G418 to indicate which spore contains the deletion and C-URA to indicate the presence of the plasmid expressing the indicated gene. **A)** *S. cerevisiae lcd1∆*::KanMX strain containing a plasmid with the *S. uvarum* allele of *LCD1.* **B)** *S. uvarum aft1∆::*KanMX strain containing a plasmid with the *S. cerevisiae* allele of *AFT1.*

## Discussion

In this study, we applied a comparative functional genomics approach to investigate how genetic background influences gene dispensability between the reference strains of two diverged species of yeast. Using insertional integration comparisons between haploid and diploid pools of mutants, we prioritized genes to validate as predicted essential, non-essential, and differentially essential gene categories in *S. uvarum.* We predicted approximately 12% of orthologs to differ in dispensability between *S. uvarum* and *S. cerevisiae* and validated 22 genes in this category. Surprisingly, however, most genes that differ in dispensability have retained their function between these two species, suggesting that differences in gene dispensability are likely due to *trans-*acting changes rather than the direct result of divergent coding sequence.

Specifically, our comparison of orthologous genes between *S. cerevisiae* and *S. uvarum* revealed that a majority of genes maintain conserved dispensability requirements (88%) while 12% of orthologs are predicted to be essential in one species but not the other. We confirmed 93% (15/16) of predicted conserved categories of essentiality and 49% (27/55) of genes predicted to be differentially essential. While applying a less restrictive insertion ratio cut-off value includes more genes to be characterized in this category, it also increases the likelihood of false positives and creates a challenge for the correct validation of this category type. Although our rate of confirmed genes in this category was lower than the conserved category, we correctly identified a subset of genes that are differentially dispensable, despite the moderately dense insertional profile of the library and a less restrictive cut-off value applied to include more genes to be classified as this type. Further analysis of predicted species-specific essential genes revealed enriched GO ontology terms of molecular functions involved in structural constituent of the ribosome and DNA binding, although more precise analysis of functional enrichments may require more thorough validation to remove the influence of false positives. Finally, we utilized yeast genetic tools to test hypotheses about genetic background effects that contribute to differences in essentiality. We find that differences can be explained by paralog divergence and *trans*-acting factors.

Applying a random insertional approach has proved to be useful in functionally profiling *S. uvarum* and will be useful for studying other understudied species, with the goal of adding information to gene annotation methods. While this study was performed in standard rich media laboratory conditions, it is easily amenable for testing stressful conditions, other nutrient sources, as well as naturally relevant conditions. This library can be applied to probe previously un-annotated genes or even proto-genes for functional acquisition, since it is not restricted to *a priori* assumptions of genic boundaries. The identification of synthetic lethal interactions can also be determined by performing insertional profiling in the background of a particular mutation of interest relatively quickly and economically. Similarly, the library could be generated in different strain backgrounds to confirm which phenotypes are truly species-specific versus which might vary within the species. Additionally, pooled competition experiments *en masse* can be used to determine the frequency of particular insertional mutants, providing quantitative measurements of cellular fitness across conditions. Such a strategy could be efficiently employed using computational approaches to prioritize experimental conditions that are most likely to probe the most valuable phenotypic information for further functional characterization (Guan et al. 2010).

Gene regulation also plays a large role in evolution and is crucial for responding to environmental change (Carroll 2005). In previous studies, we aimed to experimentally characterize differences in gene expression patterns between *S. cerevisiae* and *S. uvarum* and discovered species-specific responses to osmotic stress, peroxisome biogenesis and autophagy, suggesting that each species may have been exposed to different selective pressures within their respective evolutionary histories (Caudy et al. 2013; Guan et al. 2013). Interestingly, we did not find that genes with different gene expression patterns between species were more likely to be differentially essential. Instead, *trans* genetic interactions dominate. Identifying the molecular basis of these *trans* effects can now be undertaken, potentially revealing principles of genetic interactions across species.

## Materials and Methods

### Strains, plasmids, and primers

The strains, plasmids, and primers used in this study are listed in **Supplemental Files 1**, **2** and **3** respectively. All *S. uvarum* strains are derivatives of the sequenced strain CBS 7001 (previously sometimes called *S. bayanus* or *S. bayanus* var *uvarum*), and all *S. cerevisiae* strains are of S288C background. Unless specified below, yeast strains were grown at 25°C for *S. uvarum* strains and 30°C for *S. cerevisiae* strains in media prepared according to standard recipes.

### Construction of the Tn7 mutagenesis library

The construction of the Tn7 plasmid library has been previously described in detail and was obtained from the Caudy lab (Caudy et al. 2013). Briefly, this mutagenesis approach uses a plasmid library of *S. uvarum* genomic DNA, containing random Tn7 transposon insertions. The construct has a selectable marker for transformation into yeast, allowing the selection of disruption alleles.

To make the plasmid library, genomic DNA was isolated and fragmented by sonication to an average length of 3 kb from a ρ^0^ *S. uvarum* strain. The ends of the DNA were blunted and cloned into the pZero Blunt vector (Invitrogen). Approximately 50,000 colonies were recovered from the transformation into *E. coli* DH5α strain. The transformants were scraped from Kanamycin plates and pooled for plasmid purification. A version of the Tn7 transposon was constructed by amplifying the promoter from the Tet-on pCM224 (Bellí et al. 1998). The cassette of the Tet-on promoter and the ClonNAT resistance gene was amplified using PCR primers containing *lox* and BamHI sites and cloned into the BamHI site of the NEB vector pGPS3. This transposon construct was inserted into the *S. uvarum* genomic DNA library *in vitro* using the transposon kit from NEB. Initial selection (50,000 colonies) was on ClonNAT/Zeo. HindIII and XbaI were used to digest the pZero backbone to release the linearized genomic DNA for efficient recombination. The library was then used to transform a haploid *S. uvarum* strain (ACY12) and a diploid strain (YMD1228) using a modified transformation protocol optimized for *S. uvarum* (Caudy et al. 2013). Transformant colonies were plated to YPD-ClonNat plates and allowed to grow for 5 days at 25°C. A total of ~ 500,000 colonies were scraped for each pool. Each final pool was well mixed at a 1:1 ratio with 50% glycerol and 2 ml aliquots were stored at −80°C.

### Pooled growth of Tn7 *S. uvarum* libraries

To determine the initial complexity of the integrated pools, genomic DNA was extracted directly from the glycerol stocks of both haploid and diploid pools using the Hoffman and Winston method (Hoffman and Winston 1987). Additionally, we inoculated 500 μl of both libraries in separate YPD flasks for 24 hours to recover mutants after 24 hours of growth. Furthermore, to collect samples over time, we competed both pools under sulfate-limiting conditions in chemostats for approximately 30 generations at 25°C. A large-volume, ~300ml, sulfate-limited chemostat (Gresham et al. 2008) was inoculated with a single 2ml glycerol stock sample of each pool. After allowing the chemostat to grow at 25°C without dilution for ~24 hrs, fresh media was added to the chemostat at a rate of 0.17 h^−1^. This pooled growth assay was repeated twice, each including 5 time points with O.D. and dilution rate measurements as well as collected cell pellets for DNA extractions using the modified Hoffman-Winston prep referenced above. Early time-points from this pooled growth assay were included in our analysis here because we found it to be largely overlapping with the rich media collection, and we found that the additional sequencing coverage improved our overall results.

### Tn7 sequencing library preparation

Sequencing libraries were prepared by first extracting genomic DNA from pools of each library grown in YPD and sulfate limited conditions. Genomic DNA libraries were prepared for Illumina sequencing using a Tn7-seq protocol described previously (Wetmore et al. 2015). Briefly, the Covaris was used to randomly fragment DNA to approximately 200-800 bp in length. The fragments were blunt ended and A-tails were added to the fragments to ligate the Illumina adapter sequences. Custom index primers (listed in **Supplemental File 3**) targeting Tn7-specific sequence and Illumina adapter sequence were used to enrich for genomic DNA with Tn7 insertion sites. The barcoded libraries were quantified on an Invitrogen Qubit Fluorometer and submitted for 150 bp-paired end sequencing on an Illumina HiSeq 2000 by JGI. This method was also applied to make the plasmid library, from linearized plasmid DNA.

### Sequencing analysis

Sequencing reads from the FASTQ files were trimmed to remove Tn7 specific sequences and adapter sequences, restricting the minimal length of reads to 36 bp using Trimmomatic (Bolger et al. 2014) and FASTX-Toolkit. Trimmed FASTQ files were aligned against the reference strain of *S. uvarum* (CBS 7001) using Burrows-Wheeler Aligner (BWA) with standard filters applied (Li and Durbin 2009). Specifically, non-uniquely mapping reads, reads in which the pair did not map, reads with a mapping quality less than 30 and PCR/optical duplicate reads were filtered out; the samtools C-50 filter was applied as recommended for reads mapped with BWA. To limit the insertional analysis to actively growing cells, SAM files were merged from the later time points in the growth assays of each pool using samtools (Li et al. 2009). The sequence coverage of the nuclear genome ranged from 70 to 300x (**Supplemental File 4**). Insertion sites were determined from SAM files using a custom Ruby script. Insertion sites that had 10 reads or more were processed through a custom Python script that counted the number of insertion events in each coding region across the genome. This pipeline was applied to both libraries and further comparisons were made between the pools to determine essential genes. Read data have been deposited at the NCBI under the SRA accession number SRP115313.

### Predicting gene dispensability between species

In order to determine a list of predicted essential genes, comparisons were made between the haploid and diploid libraries. We calculated an insertion ratio by dividing the number of insertions in the haploid pool by the number in the diploid pool. This direct comparison inherently accounts for the length of the gene, since the length is constant in both libraries. Therefore, a decrease in insertion sites in the haploid library indicates a reduction in the presence of mutants containing insertional sites that impact cellular viability. Ratios closer to zero represent insertional mutants that reduce the frequency of haploids harboring insertional sites in a coding region that is required for cellular growth.

To make an insertion ratio cut-off value to categorize essential and non-essential genes, we analyzed the distribution of insertion ratios within intergenic regions between 500 bp and 7 kb in length and positioned between Watson and Crick coding regions (so chosen because these are less likely to contain promoter sequences). The distribution of the insertion ratio calculated for these regions was similar to that of known non-essential genes in *S. cerevisiae.* Therefore, we used this distribution to rank the insertion ratios of all coding regions and set a cut-off value to 0.25 where 20% of the insertion ratio of coding regions fell below the intergenic distribution, which was similar to the kernel density estimates of known *S. cerevisiae* essential genes. The kernel density estimates were computed in R and visualized using ggplot2. To remove a class of low coverage genes in the essential gene category, we applied an additional cut-off value. Since the difference between 0 and 1 with a gene that is longer has a lower weighted difference than a shorter gene, we calculated the difference between the diploid pool and haploid pool and normalized this value to the length of the gene (normalized difference). Genes with less than a normalized difference of 2 were removed from the essential category.

### Validating predicted essential and non-essential genes

We validated predicted essential genes by creating *S. uvarum* heterozygous diploid deletion mutants using primers listed in the **Supplemental File 3.** Primers containing 50 bp of homology upstream and downstream of each candidate open reading frame were used to amplify the KanMX cassette from the pRS400 plasmid. The PCR product was used to integrate into the *S. uvarum* genome using an *S. uvarum* specific transformation protocol. The proper integration of the construct was validated through clone purifying positive colonies and extracting genomic DNA to perform PCR using diagnostic primers listed in the **Supplemental File 3.** The diagnostic primers were designed to target ~150 bp upstream and ~150 bp downstream of the open reading frame to identify wild-type and drug-marker alleles. Positive clones were sporulated for 3-5 days at 25°C and 8 tetrads were screened for 2:2 viable segregation. Images were taken after 4 days of growth on YPD plates. Mutants conferring non-essential phenotypes were replicated on G418 plates and images were taken after 4 days of growth at 25°C (**Supplemental Fig. 9&10**). This method was also applied to making double mutants. A collection of 440 *MATα S. uvarum* strains was generated by standard methods in the Rine lab and used as confirmed non-essential genes.

### Cross-species complementation assays

To determine if genes are diverging in gene function or in other t*rans*-acting factors, we performed cross-species complementation assays with species-specific essential genes. Essential genes that were *S. cerevisiae* specific were tested in a heterozygous diploid deletion strain from the SGA marker collection. Alleles of each *S. cerevisiae* essential gene and their promoters were amplified from *S. cerevisiae* and *S. uvarum* genomes and cloned into a CEN ARS plasmid. PCR using Phusion DNA polymerase was used to amplify 500 bp upstream and 5 bp downstream of the stop codon of each gene from *S. cerevisiae* and *S. uvarum.* Each gene was cloned into pIL37 by Gibson assembly using primers listed in **Supplemental File 3** using standard methods (Thomas et al. 2015). All plasmids used in this study are listed in **Supplemental File 2.** The *S. cerevisiae* heterozygous diploid deletion strains were transformed with a plasmid containing a corresponding allele from each species and selected on C-URA plates. Similarly, *S. uvarum* specific essential genes were also tested by making each heterozygous diploid deletion strain *ura3∆/ura3∆,* and transformed with a plasmid containing a corresponding *S. cerevisiae* allele from the MoBY-ORF collection (Ho et al. 2009).

Transformed strains were sporulated for 5 days at 30°C and 25°C for *S. cerevisiae* and *S. uvarum* species, respectively, and tetrad analysis was performed on YPD plates. After 3 days of growth, plates were replica plated on C-URA and YPD+G418 plates and imaged after 2 days of growth (**Supplemental Fig. 15**).

### Data Access

The transposon insertion data in this study have been submitted to the NCBI Sequence Read Archive (SRA; https://www.ncbi.nlm.nih.gov/sra/) under SRA accession number SRP115313.

## Acknowledgments

Thank you to Jasper Rine for generously sharing *S. uvarum* deletion strains. Thank you to Noah Hanson and Kolena Dang for technical assistance with tetrad dissections. Thank you to Daniel Chee for his help with optimizing the Python code and Jeremy Stone for his help with the Ruby script that was used for the insertional analysis pipeline. We thank M.K. Raghuraman and Harmit Malik for their helpful comments on the manuscript and thank Sergey Kryazhimskiy for his helpful insight about data analysis. This work was supported by NSF grant 1516330. MS was funded by NSF GSRF DGE-1256082 and the Robert D. Watkins Graduate Research Fellowship from the American Society for Microbiology. MJD is supported in part by a Faculty Scholars grant from the Howard Hughes Medical Institute. MJD is also a senior fellow in the Genetic Networks program at the Canadian Institute for Advanced Research. Sequencing resources were provided by JGI CSP Project 1460. AAC was supported by an Open Operating Grant from the Canadian Institutes for Health Research and by a Discovery Grant from the Natural Sciences and Engineering Research Council of Canada. AAC is the Canada Research Chair of Metabolomics for Enzyme Discovery.

## Author contributions

**Conceptualization:** MRS CP AC MJD.

**Formal analysis:** MRS MJD.

**Funding acquisition:** AC MJD APA RB.

**Investigation:** MRS CP FC BH MJD.

**Project administration:** MRS CP FC BH MJD.

**Resources:** JMS RB APA AC.

**Supervision:** MJD.

**Visualization:** MRS.

**Writing – original draft:** MRS MJD.

